# Role of the Nucleotide Excision Repair endonuclease XPF in the kinetoplastid parasite *Trypanosoma brucei*

**DOI:** 10.1101/2025.01.22.633349

**Authors:** Claudia Gómez-Liñán, María Sáez-Maldonado, Luís Miguel Ruíz-Pérez, Dolores González-Pacanowska, Antonio E. Vidal

## Abstract

The nucleotide excision repair (NER) mechanism is responsible for the removal of bulky DNA damage such as pyrimidine dimers induced by ultraviolet light. The NER pathway detects such lesions and excises the damaged strand through incisions at 5 ’ and 3’ of the damage. The 5’ incision is catalyzed by a heterodimeric endonuclease composed of XPF (catalytic subunit) and ERCC1 (non-catalytic). Here, we show that the genome of *Trypanosoma brucei*, the causal agent of human African trypanosomiasis or sleeping sickness, codes for an XPF ortholog. RNAi silencing of TbXPF sensitizes cells to UV irradiation, thus providing evidence that NER operates in these parasites. In addition, TbXPF confers protection against intra- and inter-strand crosslinks induced by cisplatin and mitomycin C respectively. Consistent with a role in DNA repair, XPF localizes to the cell nucleus, and is found associated to nucleoplasmic and nucleolar regions. The presence of a functional NER pathway in trypanosomes suggests that *in vivo*, they are susceptible to undergo replication and transcription-blocking DNA damages. The results obtained with various antitumor agents provide proof of concept for the potential of NER inhibition as a means to improve antiparasitic therapies.

## INTRODUCTION

Nucleotide excision repair is a versatile DNA repair system responsible for the removal of diverse helix-distorting bulky lesions such as those formed by UV light, environmental mutagens, and some cancer chemotherapeutic adducts. In humans, defective NER leads to several autosomal recessive disorders such as xeroderma pigmentosum (XP), Cockayne’s syndrome (CS), UV sensitive (UV^s^) syndrome (UVSS), cerebrooculofacioskeletal (COFS) syndrome and trichothiodystrophy (TTD) ^1^. The mammalian NER pathway uses more than 30 proteins to detect the DNA lesion, remove around 24-32 nucleotides of the damaged strand, fill the gap by using the undamaged strand as a template and complete the error-free repair by sealing the strand ^2^. In NER, two damage detection mechanisms exist, global genome repair (GG-NER) that eliminates DNA lesions throughout the genome, and transcription-coupled repair (TC-NER) that specifically repairs damage from the template DNA strands of actively transcribed genes. Thus, while GG-NER is initiated by the recognition of DNA helix-distorting damage, a blocked RNA polymerase II (RNA Pol II) at a lesion constitutes the first step for damage recognition in TCR. The XPC complex formed by XPC, RAD23B and centrin 2 and the UV–DDB (ultraviolet radiation–DNA damage-binding protein) complex, consisting of DDB1 and DDB2, constitute the main DNA damage sensors in GG-NER. In TC-NER, lesion-stalling of RNA Pol II recruits the TC-NER-specific factors Cockayne syndrome proteins CSA and CSB, required for further assembly of the NER machinery. Once the lesion is detected, both subpathways converge into the same route. First, DNA damage is verified by TFIIH (transcription initiation factor IIH), which is a transcription initiation and repair complex containing ten protein subunits including XPB and XPD helicases. The existence of a DNA lesion is further verified by the XPA protein, which binds to single-stranded, chemically altered nucleotides. The next step is DNA damage excision, catalyzed by the structure-specific endonucleases XPF–ERCC1 (recruited by XPA) and XPG, which incise the damaged strand at 5’ and 3’ from the lesion, respectively. Gap-filling DNA synthesis starts right after 5’ incision and involves PCNA, RFC, RPA and the use of distinct DNA polymerases and ligases depends on the cells proliferation status. DNA synthesis by DNA Pol ε and sealing by DNA ligase 1 occurs in replicating cells while DNA Pol δ, DNA Pol κ and XRCC1-DNA ligase 3 act in non-replicating cells (reviewed in ^2,3^).

Both XPF (ERCC4; excision repair cross-complementing group 4 protein) and ERCC1 belong to the XPF/MUS81 family ^4^. They form a stable heterodimer composed of catalytic (XPF) and non-catalytic (ERCC1) subunits. The XPF–ERCC1 complex (Rad1–Rad10 in yeast) cleaves DNA at the 5′ of the UV-induced lesions by recognizing the presence of 3′ ssDNA at a double-strand/single-strand DNA junction ^5^. XPF– ERCC1 performs additional DNA repair functions outside of NER and is involved in double strand break repair, base excision repair, DNA-DNA crosslink repair and maintenance of telomeres ^6–9^. The multifunctional role of the complex XPF-ERCC1 underlies the remarkable array of disorders found in human patients and mouse models defective in XPF or ERCC1 ^10^. Thus, mutations in *XPF* and *ERCC1* genes have been associated to a variety of clinical phenotypes including abnormal skin photosensitivity, late onset of skin cancers and of neurological disease, and accelerated aging in both human patients and mice ^11^.

The unicellular protozoan parasite *Trypanosoma brucei* is the causative agent of African Trypanosomiasis (HAT) or sleeping sickness in humans and nagana disease in cattle. These parasites are transmitted through the bite of a tsetse fly (Glossina spp.). Once in the mammalian host, trypanosomes replicate extracellularly in the bloodstream until they cross the blood–brain barrier to initiate the second stage of the disease, which causes the characteristic sleep disorder ^12^. HAT affects million of people throughout sub-Saharan Africa and is usually fatal if untreated or inadequately treated. According to the World Health Organization, HAT remains one of the most important and challenging neglected tropical diseases to eradicate due, among other factors, to the need of new therapeutic strategies ^13^.

DNA repair is one of several mechanisms by which a cell maintains the integrity of DNA, thus ensuring the faithful transmission of its genetic material and the survival of the species. The DNA damage repair pathways are highly conserved in eukaryotes and in spite of that, perform specialized functions ^14^. Many of the GG-NER and TC-NER mammalian components have been identified in *T. brucei* ^14^, yet others seem to be absent (e.g. XPA and DDB2), perform functions outside of NER (e.g. XPC, XPD) ^15^ or are present in the genome as paralogs (e.g. XPBz and XPB) ^16^. Reverse genetics analyses have shown the implication of TbCSB, TbXPBz and TbXPG but not of other NER factors such as TbXPB, TbXPD or TbXPC, in protection against UV-induced DNA damage and/or cisplatin DNA adducts ^15,16^. Altogether, these findings suggest major divergences relative to NER in other eukaryotes, most notably, the apparent exclusion from NER of XPA and the multisubunit transcription factor TFIIH, whose core complex includes TbXPB and TbXPD but not XPBz ^17,18^. Furthermore, XPC and DDB factors are essential for cell viability and do not seem to act as DNA damage sensors, which argues against the existence of GG-NER in trypanosomes. Instead TC-NER would be the preferred NER pathway and maybe the only one operating in *T. brucei*, a choice that is likely influenced by the extent and predominance of RNA polymerase II polycistronic transcription in kinetoplastids ^15^.

Here, we have characterized the *T. brucei* ortholog of XPF (TbXPF), an essential component of the NER pathway. We present evidences of the presence of a functional NER pathway in trypanosomes, which implies that *in vivo*, these parasites undergo replication and transcription-blocking DNA damage that needs to be eliminated. Furthermore, XPF-deficient cells are highly sensitive to various antitumor agents thus providing proof of concept for the potential of NER inhibition as a means to improve antiparasitic therapies.

## RESULTS

### XPF-ERCC1 from *Trypanosoma brucei*: role in the elimination of UV-induced DNA damage

The exposure of cells to UVC radiation leads to different types of DNA damage, most of them dipyrimidinic photolesions, cyclobutane pyrimidine dimers (CPDs), and pyrimidine-(6,4)-pyrimidone products (6-4PPs), which are primarily repair by the NER pathway. While NER can be initiated either by GGR (global genome repair) or TCR (transcription-coupled repair), the final steps are common for both sub-pathways and involve among other activities, the incision at the 5′ side of the DNA damage by the nuclease XPF-ERCC1. XPF and ERCC1 form a stable heterodimeric complex that is essential for NER. Using the homology detection tool BLAST and the human proteins as the query sequences, XPF (Tb927.5.3670) and ERCC1 (Tb927.7.2060) genes were identified in the *T. brucei brucei* TREU927 genome database (TritrypDB). Pairwise alignment using the EMBOSS Needle tool (EMBL’s European Bioinformatics Institute) revealed moderate but significant homology between TbXPF and its human ortholog (identity=20.1%; similarity=31.7%). TbXPF is a protein of 1242 amino acids (136 kDa) with a well-conserved ERCC4 nuclease domain (Fig. 1A), including the metal-binding residues (GDXn(V/I/L)ERKx3D motif) that defines the active members of this family of endonucleases (Fig. 1B) ^19^. The tandem helix hairpin helix (HhH)2 domain, which in human XPF dimerizes with the equivalent (HhH)2 domain of ERCC1 to form a functional nuclease, is also present in TbXPF (Fig. 1C).

**Figure 1.**
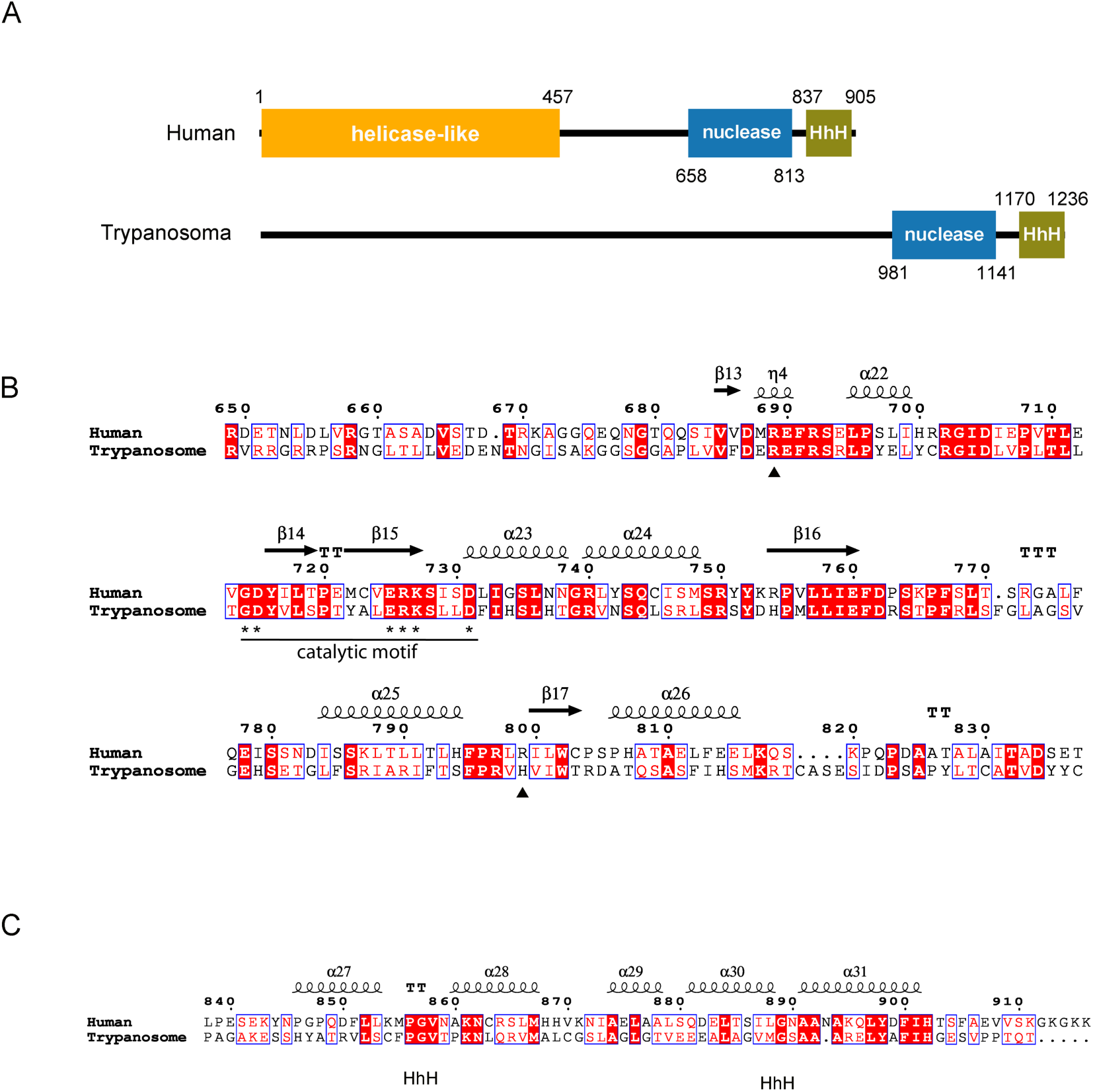
Comparative analysis of human and trypanosomal XPF proteins. (A) Schematic representation of human and trypanosomal XPF proteins. Human domains were represented according to UniProtKB/SwissProt (Q92889.3). Human XPF has a helicase-like domain that lacks key residues essential for ATP binding and hydrolysis activity ^36^. *T. brucei* protein domains were identified using the InterPro protein signature databases. No significant similarity to the human helicase-like domain was found in TbXPF (B) Sequence alignment of *T. brucei* and human XPF nuclease domains. The XPF nuclease domain extends from amino acid 658 to 813 of human XPF. Residues in this domain involved in catalysis (*) and those associated with clinical phenotypes (&) are indicated. Mutation R689S was found in patient FA104 associated to Fanconi anemia disorder; patient XP42RO with *Xeroderma pigmentosum* was homozygous for mutation R799W ^11^. (C) Sequence alignment of helix hairpin helix (HhH) domains. Sequence alignment was generated using MultAlin ^58^ and the final format was obtained with ESPript ^59^. XPF protein sequences were retrieved from NCBI RefSeq: *Homo sapiens* (NP_005227.1); *Trypanosoma brucei* (XP_845037.1).

In order to examine the role of the NER pathway in trypanosomes, an inducible stable RNAi cell line expressing a dsRNA against the ORF sequence of XPF was established (RNAi-ORF cell line). RNAi-ORF cells exhibited a 50% decrease in their *XPF* mRNA levels without affecting normal proliferation (Fig. 2A). XPF-silenced cells were, however, more sensitive to UV irradiation than uninduced and parental cells (Fig 2B). The impact of XPF on cell proliferation after UV exposure was assessed by monitoring the cell cycle progression along 12 hours. It is well established that bulky UV lesions block replicative DNA polymerases during the S phase of the cell cycle. Accordingly, an increase of cells in S-phase became evident at 3 hours post-irradiation for both repair-proficient and repair-deficient cells (Fig. 2C). However, while XPF-proficient cells started to recovered normal S-phase levels at 6 and 12 hours post-irradiation, the progression of XPF-deficient cells through the S-phase was slowed significantly over at least 12 hours. The intra-S delay observed in the absence of XPF likely reflects the persistence of replication fork-blocking lesions and additional unresolved intermediates. The proportion of cells with DNA content lower than G1 and higher than post-G2/M did not exceeded 10% in any experimental condition and the absence of XPF did not cause any significant difference.

**Figure 2.**
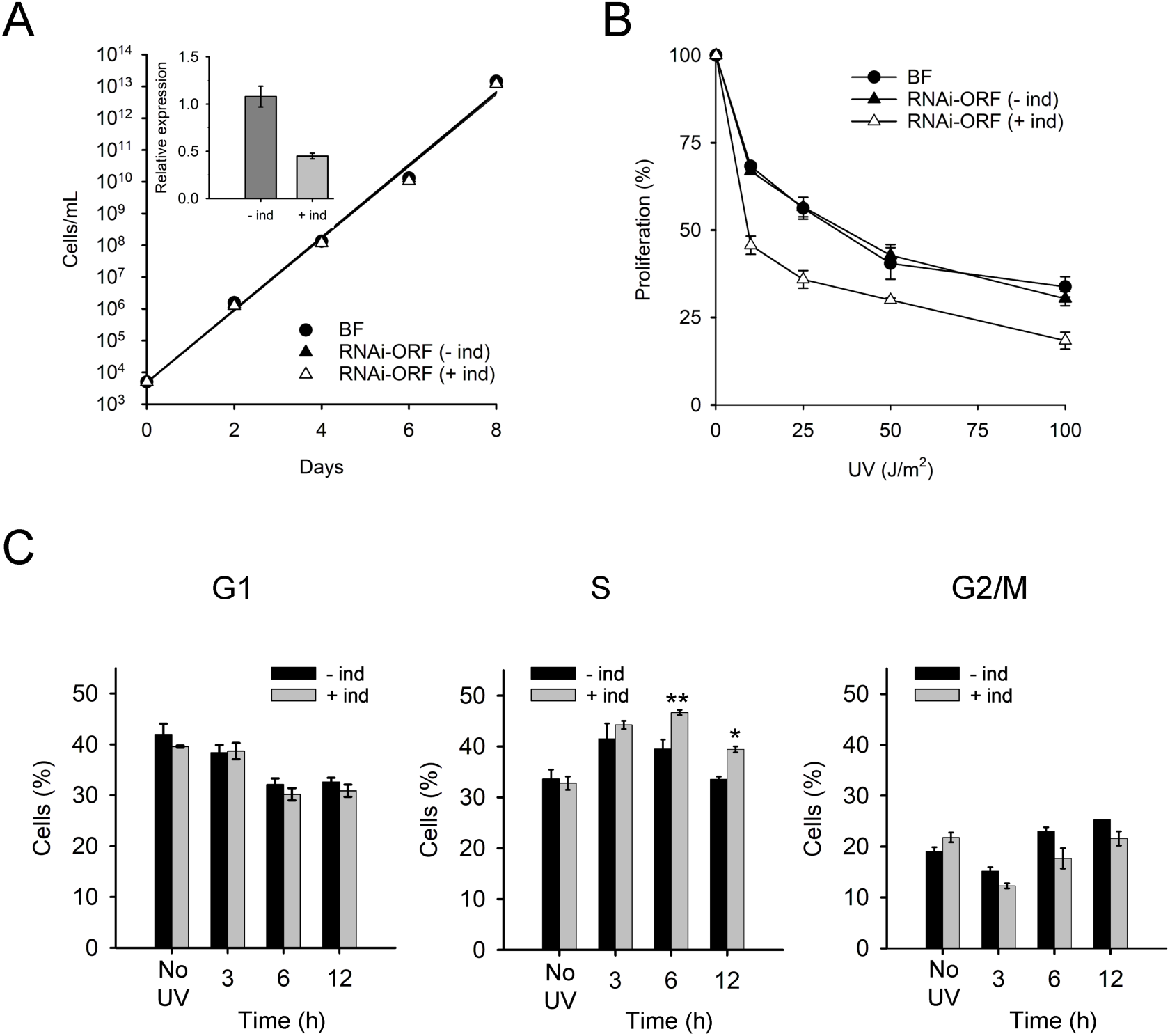
NER factor XPF protects *T. brucei* from UV photolesions. (A) Cumulative cell growth of trypanosomes cultured in standard HMI-9 medium over an eight-day period. RT-qPCR quantification of the levels of mRNA for the RNAi-targeted gene in the absence (-ind) or the presence of doxycycline (+ind), relative to mRNA levels in parental bloodstream parasites; vertical lines denote standard deviation from two experiments. Data represent the mean for two independent biological replicates. (B) UV sensitivity assay. Log-phase parasites at 5 x 10^3^ cells/mL were exposed to increasing doses of UVC irradiation and incubated for 48 h at 37 °C in HMI-9 medium before counting. Proliferation was calculated relative to cell growth in non-irradiated cells. Experiments were performed three times, values are the mean (±SD). Cell lines are represented with the following symbols: BSF (*circle*), RNAi-ORF (filled triangle), RNAi-ORF in the presence of 1 μg/mL of doxycycline (empty triangle). (C) Effect of UVC irradiation and XPF depletion on cell cycle progression. Two days after RNAi induction, XPF-depleted cells were exposed to UV light (50 J/m^2^) and samples collected at different times after irradiation. Cell cycle phase was determined by analyzing DNA content by flow cytometry (FACS). Plots show the percentages of cells for each time point in each of the different cell cycle stages: G1, S and G2/M. Represented data are the mean (±SD) of two independent determinations. *\** and *\*\** indicate P-values (t-test) of < 0.05 and < 0.01 respectively.

### XPF repairs intra- and inter-strand DNA crosslinks

In order to gain insight into the cellular function of the TbXPF, RNAi depleted-cells were exposed to several DNA damaging agents some of which are effective chemotherapy drugs used in the treatment of several cancer types. Cisplatin (CDDP) is a front-line drug in the treatment of lung, colorectal, ovarian, and head-and-neck cancers. In human cells, cisplatin-induced DNA lesions, mostly intra-strand diadducts ^20^, are repaired primarily by nucleotide excision repair (NER) ^21,22^. On the other hand, mitomycin C (MMC) is classified as an alkylating agent capable of covalently binding DNA and inducing inter-strand cross-linked lesions ^23^. Inter-strand crosslinks (ICLs) exert their cytotoxic action by interfering with DNA replication and transcription ^24^. As shown in Fig 3A and 3B, both, CDDP or MMC strongly impaired parasite proliferation, an effect that was much more pronounced in parasites with reduced levels of XPF. These results suggest a role in intra- and inter-strand DNA crosslink repair for the trypanosomal XPF endonuclease.

**Figure 3.**
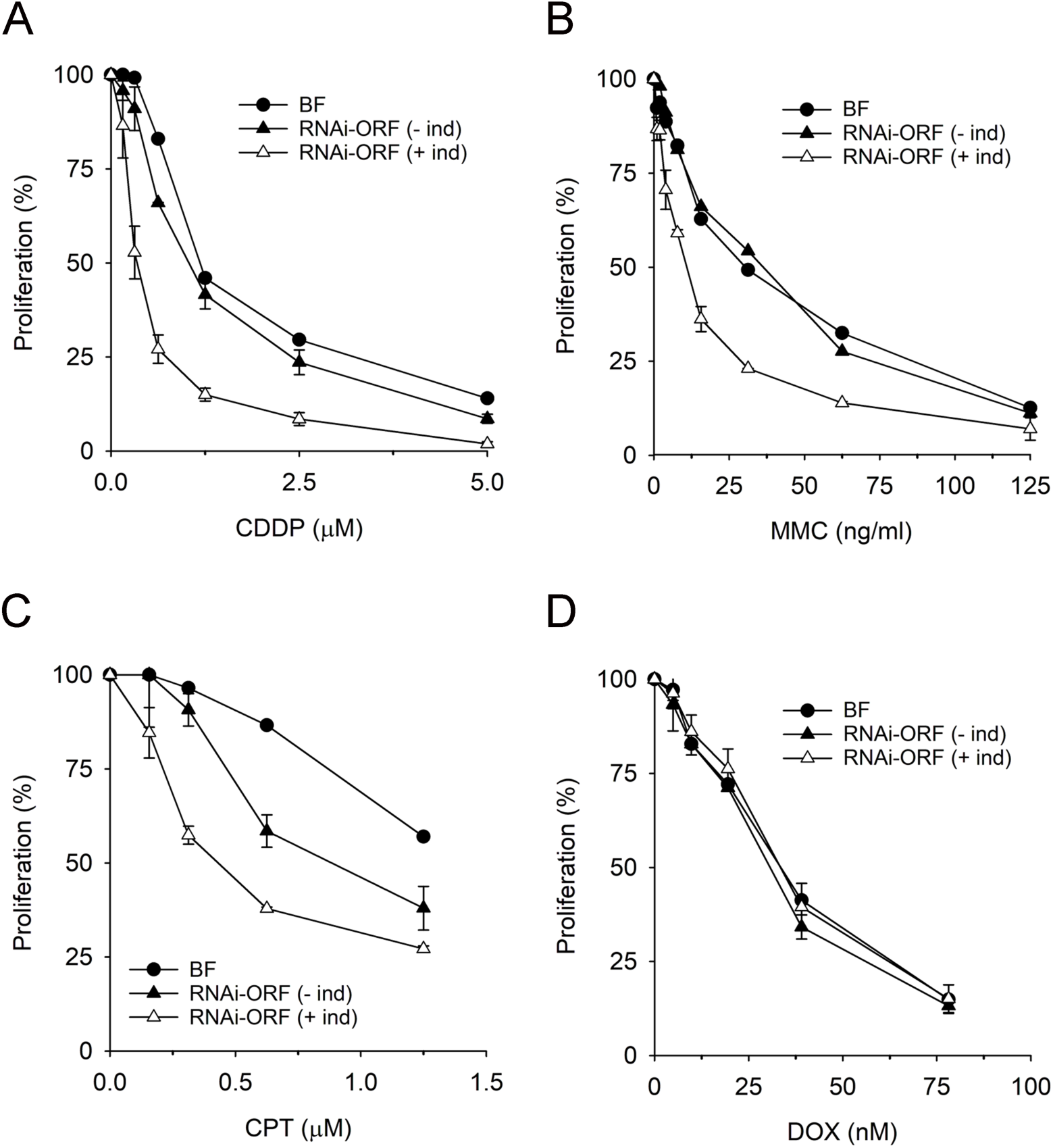
XPF-deficient cells exhibit increased sensitivity to anticancer agents. Log-phase parasites at 5 x 10^3^ cells/mL were exposed to increasing concentrations of (A) cisplatin (CDDP); (B) mitomycin C (MMC); (C) camptothecin (CPT); (D) doxorubicin (DOX) for 24 h. at 37 °C. Cell growth was measured with resazurin as described in Methods. For each cell line, proliferation was calculated relative to cell growth in the absence of compound. Experiments were performed at least three times, in duplicate. Values are the mean (±SD).

The role of XPF was further investigated by exposing the cells to camptothecin (CPT) and doxorubicin (DOX), both considered Topoisomerase poisons that ultimately generate DNA-protein crosslinks. CPT blocks the reaction catalyzed by DNA Topoisomerase I (Topo I), resulting in the accumulation of a covalent complex between the enzyme and the 3′ end of the cleaved DNA ^25^. Conversely, doxorubicin intercalates into unreplicated DNA and inhibits of topoisomerase II activity, thus blocking DNA unwinding during transcription ^26^. Unlike Topo I-DNA crosslinks, Topo II-DNA covalent complexes would involve phosphotyrosyl linkages with a 5’ DNA end. Both, CPT and DOX are highly toxic to trypanosome cells (Fig. 3C and 3D). Down- regulation of XPF strongly increased the sensitivity to CPT whereas treatment of trypanosome cells with increasing concentrations of DOX did not reveal significant differences in XPF-silenced cells. These observations suggest that XPF participates in the repair of 3’-DNA-Topo I crosslinks or other genotoxic intermediates that result from them.

We have also investigated the contribution of NER to fexinidazole (FEX) resistance. FEX, a 5-nitroimidazol derivative, is the first all-oral drug for gambiense sleeping sickness. It has been proposed that FEX nitroreduction results in the formation of reactive chemical species with potential to react with DNA and cause genetic damage^27^. As shown in Suppl. Fig. 1A, XPF does not confer genotoxic resistance to FEX thus ruling out the generation of NER substrates by this drug as part of its mechanism of action. Other types of DNA damage such as single-strand DNA breaks and oxidative base damage were also not DNA repair substrates for XPF as RNAi-silenced parasites behaved identical to parental cells when exposed to phleomycin or hydrogen peroxide (Suppl. Fig. 1B and 1C).

### Subcellular localization of TbXPF

The subcellular distribution of the trypanosomal XPF ortholog was determined using a transgenic cell line that expresses an inducible myc-tagged version of TbXPF in cells where the RNAi targets the 3’-UTR of XPF (RNAi-UTR cell line). Upon induction, endogenous *XPF* mRNAs are depleted while ectopic myc-TbXPF, which lacks the UTR sequences, escapes silencing (Fig. 4A). As expected, RNAi-UTR induced cells exhibit the same UV-sensitive phenotype observed in RNAi-ORF cells while the expression of myc-TbXPF restores normal proliferation (Fig 4B). The phenotypic rescue by myc- TbXPF validates the myc-tagged version as a functional protein suitable for subcellular localization studies. Indeed, staining with anti-myc antibodies showed that XPF is present in the nuclear compartment (Fig. 4C), a localization that is consistent with its DNA repair function. A closer inspection revealed XPF expression in several nucleoplasmic and nucleolar foci or dots (upper row). In those cases where only one dot was detected in the nucleus, it was exclusively present in the nucleolus (lower row). Similar nuclear distribution was observed using two different primary antibodies (Fig. 4C and Suppl. Fig. 2). Nucleolar localization was confirmed by double-staining of XPF and L1C6, a known nucleolus marker ^28^, which showed both proteins in close proximity in the nucleolar region. Although it is unclear its functional significance, various DNA repair proteins have been found associated to the mammalian nucleoli. It remains to be determined whether the nucleolus merely acts as a storage site for DNA damage response proteins or if these proteins perform specific nucleolar roles ^29^,

**Figure 4.**
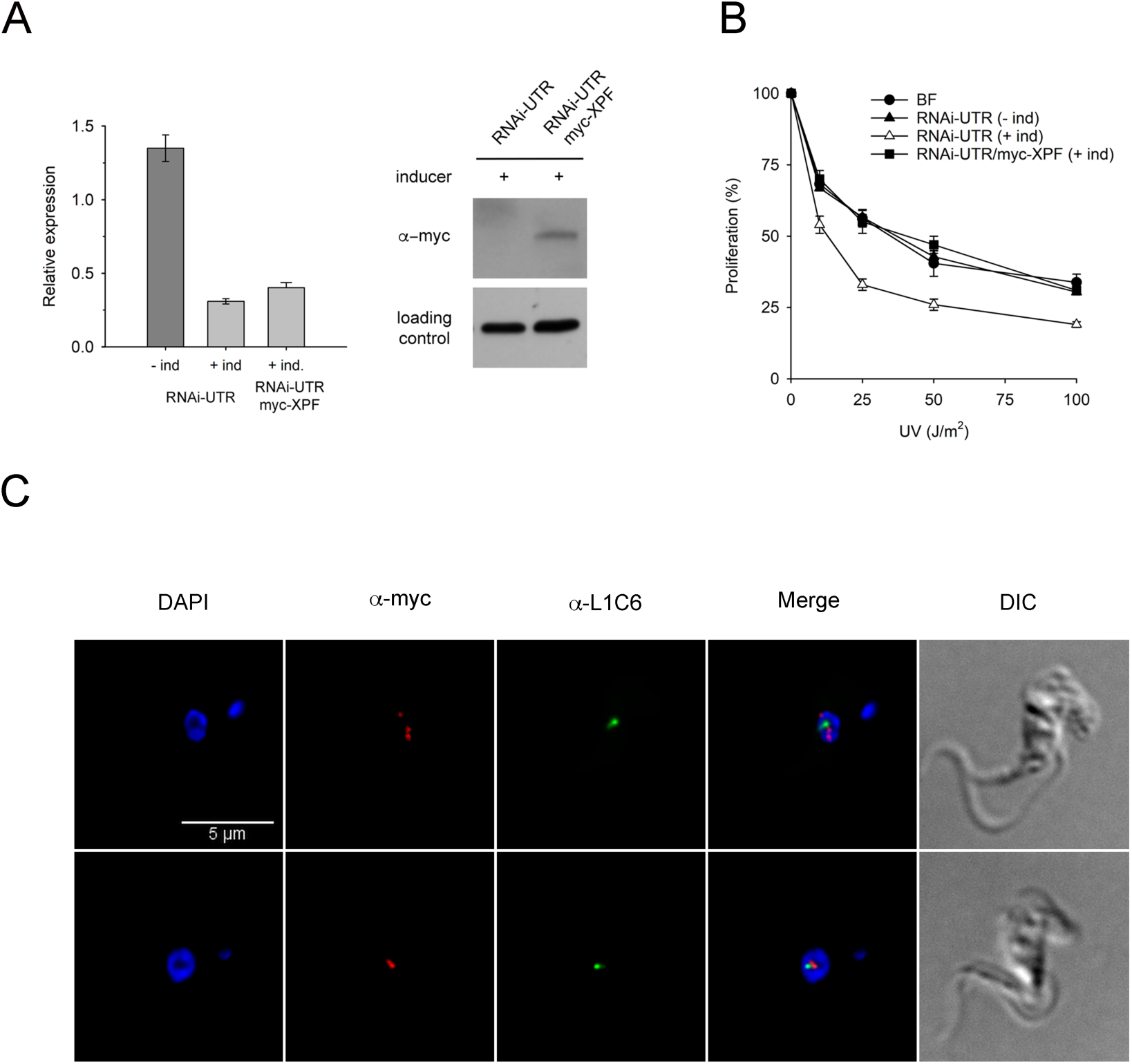
XPF localizes to the nuclear and nucleolar foci in bloodstream *T. brucei*. (A) Left, levels of mRNA were determined by RT-qPCR for each cell line and calculated relative to mRNA levels in parental bloodstream form; vertical lines denote standard deviation from two experiments. Right panel, western blot showing myc- TbXPF protein levels in whole cell extracts from 5 x 10^6^ parasites. Myc-TbXPF was detected with the anti-myc monoclonal antibodies. As loading control, ITPA protein was measured with anti-ITPA polyclonal antibodies ^60^. Images were cropped and aligned for easier comparison. (B) Log-phase parasites at 5 x 10^3^ cells/mL were exposed to UVC irradiation and incubated for 48 h at 37 °C in HMI-9 medium before counting. RNAi and protein expression were pre-induced by adding doxycycline (1 μg/mL) to the corresponding cell lines 48 h before to the experiment. Proliferation was calculated relative to cell growth in non-irradiated cells. Experiments were performed three times, values are the mean (±SD). Cell lines are represented with the following symbols: BSF (circle), RNAi-UTR (filled triangle), RNAi-UTR +ind (empty triangle), RNAi-UTR/myc-XPF +ind (filled square). (C) Immunofluorescence microscopy images were obtained to determine the subcellular localization of mycTbXPF in overexpressing cells. Nuclear and kinetoplast DNA were stained with DAPI. Myc-XPF was visualized with anti-myc tag rabbit monoclonal antibody (clone 71D10) and Alexa Fluor 594 goat anti-rabbit. Nucleolar protein L1C6 was detected with anti-L1C6 mouse monoclonal antibody and Alexa Fluor 488 goat anti-mouse. Images were collected with an inverted Leica DMi8 microscope, 100x objective, and LASX software.

### N3-Alkylation DNA damage is repaired by NER in trypanosomes

Monofunctional SN2-alkylating agents such as MMS (methyl methanesulfonate) induce primarily the N-alkylation of purine bases in DNA. N3-methyladenine (N3-meA) adducts are the primary cytotoxic lesion induced by MMS and if not repaired they can block DNA replication and cause further DNA damage and eventually cell death ^30^. N3- meA is usually repaired by the Base Excision Repair (BER) pathway, which is initiated by a methyladenine DNA glycosylase that excises the damaged DNA base lesion ^1^. Accordingly, prokaryotic and eukaryotic cells deficient in AP endonuclease activity have an increased sensitivity to the toxic effects of MMS due to their inability to repair alkylation damage via BER ^31–33^. On the other hand, O6-position of guanine represents a major site of methylation by SN1-type alkylating agents like MNNG (methyl-nitro- nitrosoguanidine). While O-methylguanine (O6-meG) lesions are produced in a lower proportion than N-methyl adducts, they are the cause of many of the cytotoxic biological effects of alkylating agents. O6-meG adducts are reverted to guanine by the action of a methylguanine methyltransferase protein ^34^.

In order to determine the cytotoxic impact in *T. brucei* of the two main genotoxic lesions induced by methylating agents, N3-meA and O6-meG, the sensitivity of the parasite to MMS and MNNG was anlyzed. The specfic impact of each DNA lesion can be inferred from the cellular response to these agents in the presence of methyladenine glycosylase 1 from *Saccharomyces cerevisiae* (MAG1) that recognizes and repairs N3- meA or human methylguanine methyltransferase (MGMT), which repairs specifically the O6-meG lesion. As shown in Suppl. Fig. 3A, impaired cell proliferation induced by MMS can be reverted by the overexpression of MAG1 and to a lesser extent of MGMT, indicating that both, N3-meA and O6-meG base damages are responsible for the cytotoxic effect. In contrast, only MGMT can revert the sensitivity to MNNG that in this case can be fully attributed to formation of O6-meG adducts.

We previously reported that *T. brucei* parasites deficient in AP endonuclease activity do not exhibit an increased sensitivity to the toxic effects of MMS ^35^. To investigate whether parasites use NER instead of BER for the removal of methyl DNA adducts, XPF-deficient cells were treated with MMS and MNNG (Fig. 5). Cell proliferation curves show that XPF depletion increases the sensitivity to MMS indicating that N3- methyl DNA lesions are indeed repaired by the NER pathway (Fig. 5A). On the contrary, XPF does not confer resistance to MNNG and therefore O6-meG lesions are not physiological substrates for trypanosomal NER (Fig. 5B).

**Figure 5.**
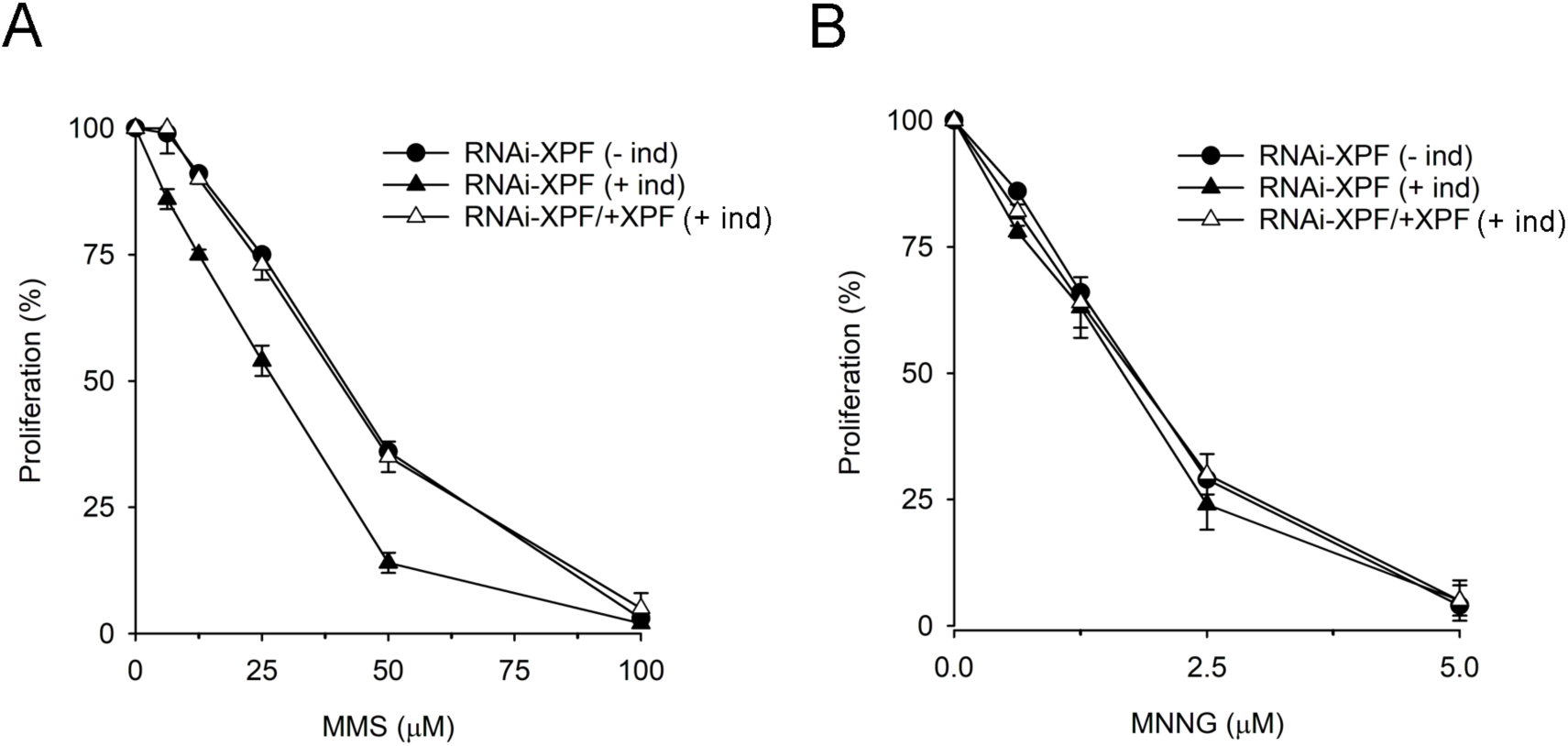
Trypanosomal NER is involved in the repair of N3-methylA adducts. Log-phase parasites at 5 x 10^3^ cells/mL were exposed to increasing concentrations of (A) methyl methanesulfonate (MMS) and (B) methyl-N-nitrosoguanidine (MNNG) for 48 h at 37 °C before counting. For each cell line, proliferation was calculated relative to cell growth in the absence of compound. Experiments were performed at least three times, in duplicate. Values are the mean (±SD).

## DISCUSSION

Nucleotide excision repair (NER) is the main pathway used by prokaryotic and eukaryotic organisms to remove a wide range of bulky DNA lesions such as those formed by UV light. The presence of most of the eukaryotic NER genes in the genome of kinetoplastid parasites suggests the conservation of practically all NER functions, including the strategies for DNA damage detection that define global-genome and transcription-coupled repair subpathways. However, early studies have shown a specialization of these parasites at transcription-coupled repair, highlighting the functional divergence of many of the trypanosomal NER factors ^15,16^.

XPF-ERCC1 is a heterodimeric endonuclease composed of two paralog subunits believed to have evolved as a result of gene duplication in lower eukaryotes ^36^. Both subunits interact through their C-terminal helix-hairpin-helix domains ^37^. This interaction is required not only for the NER function but also to preserve protein stability *in vivo* ^38^. Both, *T. brucei* XPF and ERCC1 harbor the C-terminal HhH domains which suggests that the heterodimeric form might be the true physiological state of these proteins in trypanosomes. In addition, like its human counterpart, TbXPF harbors a functional nuclease catalytic domain; all critical residues involved in DNA/metal binding and nuclease activity are present ^19^. However, the helicase-like domain of human XPF, which may also have a role in DNA binding and the subsequent cleavage activity ^36,39^, is weakly conserved in the trypanosome enzyme. An interaction between the helicase-like domain and SLX4 (Fanconi anemia complementation group P; FANCP) seems to mediate the recruitment and the specific role of the XPF-ERCC1 endonuclease in inter-strand crosslink (ICL) repair ^40^. As trypanosomes are devoid of most of the Fanconi anemia factors including an SLX4 homolog ^41^, the helicase-like domain of TbXPF may have evolved to perform its ICL repair function by establishing different protein-protein interactions yet to discover.

SLX4 is not the only XPF-ERCC1 interacting protein missing in *T. brucei*. XPA is a central factor in the coordination of NER through the interaction with almost all NER proteins including TFIIH, RPA, PCNA, XPC, DDB2 and ERCC1-XPF ^3^. Human ERCC1 interacts with XPA through its central domain and facilitates the recruitment of the endonuclease to the 5′ junction at photoproducts during NER ^39,42^. The absence of XPA in trypanosomes raises the question as to how NER is coordinated in this parasites and specifically, how the complex XPF-ERCC1 is targeted to the DNA damage site. It is plausible that a protein structurally unrelated to XPA, yet exhibiting analogous properties, could take on the role of XPA during NER ^43^.

The heterodimer formed by XPF and ERCC1 is a highly conserved protein complex, which has been identified and characterized in different model organisms. For instance, yeast *S. cerevisiae* Rad1-Rad10 null mutants are viable ^44^ and the activity of XPF- ERCC1 does not seem to be essential either in plants, worms or flies where viable *RAD1* mutants have been obtained and characterized ^45–47^. In contrast, the animal models exhibit far more severe phenotypes. ERCC1 and XPF-deficient mice are not viable or undergo early postnatal death depending on the genetic background while in humans, mutations affecting these genes have been linked to some severe diseases, including skin cancer-prone disease *Xeroderma pigmentosum* (XP), a progeroid syndrome of accelerated aging (XFE), and a cerebrooculo-facio-skeletal syndrome (COFS) ^10^.

Similarly to other lower eukaryotes, strong down-regulation of *XPF* mRNA levels did not have an apparent impact on *T. brucei* viability nor cell proliferation. This observation is in agreement with a high-throughput phenotypic screen study using RNAi target sequencing (RIT-seq) which reported that XPF depletion does not impair significantly the growth of bloodstream or procyclic trypanosomes ^48^. As XPF is known to be involved in DNA repair, any phenotypic change should be manifested under DNA damaging stress. Indeed, the *T. brucei* XPF-defective cells present a strong increase in sensitivity to UV irradiation. This observation is consistent with previous studies in yeast and mice ^10,49^ and validate the requirement of the XPF-ERCC1 complex in the NER pathway in trypanosomes.

In addition to UV-induced DNA damage, XPF confers protection from inter-strand DNA crosslinks (ICL) and TopoI-DNA complexes indicating that besides NER, it appears to be involved in other DNA repair processes. Both human XPF–ERCC1 and yeast Rad1–Rad10 have been proposed to incise 5′ to the ICL lesion and initiate the unhooking step of ICL repair. ERCC1–XPF has been shown to be required for both S-phase-dependent and –independent ICL repair ^9^. They also have been shown to be involved in repairing topoisomerase I-induced DNA damage as an alternative pathway to the single-strand break repair enzyme tyrosyl-DNA phosphodiesterase 1 (Tdp1) ^50,51^. Tdp1 processes Topo I-DNA adducts by cleaving the phosphodiester bond between the tyrosine of topoisomerase and the 3′-phosphate of DNA ^52^. In yeast, inactivation of Tdp1 has little impact on CPT cytotoxic action except in the Rad1–Rad10 null genetic background ^50,51^. Kinetoplastid genomes code for Tdp1 repair enzymes but few studies are available on them with the exception of the Tdp1 enzyme in the parasite *Leishmania donovani* ^53^. Further studies are required to determine whether XPF-ERCC1 and Tdp1 functions are also redundant in trypanosomes.

RNAi-mediated silencing of the *T. brucei XPF* gene, which codes for an essential component of the NER pathway, sensitizes cells to UV-induced DNA damage thus providing evidence that NER operates in *T. brucei*. The presence of a functional NER pathway in trypanosomes suggests that this parasite is susceptible to undergo replication and transcription-blocking DNA damage *in vivo*. It is conceivable that under genotoxic stress, genomic stability and protozoan survival may greatly depend on DNA repair systems such as NER, which for that reason could be an effective target for chemotherapeutic interventions. Further investigation will help to elucidate the exact nature of the physiological NER substrates and to determine whether the XPF function is essential in the mammalian host where the accumulation of unrepaired DNA damage may represent a challenge for parasite survival.

## METHODS

### Growth and generation of *T. brucei* cell lines

All cell lines used in this work derive from the *Trypanosoma brucei brucei* single marker bloodstream form (BF) ^54^. Bloodstream cells were cultured at 37 °C and 5% CO_2_ in HMI-9 medium supplemented with 10% (v/v) of fetal bovine serum (FBS). To generate RNAi and overexpression constructs, DNA fragments were amplified by PCR using *T*. *brucei* 427 genomic DNA as template and primers with sequences obtained from TriTrypDB (Tb427_050043200, ERCC4 domain containing protein). MAG1 open reading frame was amplified from pGAD424-MAG1, a gift from Dr. McIntyre (Institute of Biochemistry and Biophysics, Polish Academy of Sciences, Poland) while MGMT was amplified from human cDNA. Both MAG1 and MGMT were cloned into pGRV23-myc expression vector ^35,55^. All primers used in this work are listed in Supplemental Table 1. To inhibit XPF expression by RNA interference, two genetic constructs targeting either the coding or the 3’-UTR regions of XPF were generated. In these plasmids the target DNA sequences of 500-600 bp approximately, were cloned in sense and antisense directions flanking a stuffer fragment. The stem-loop cassette was produced and integrated into pGR19 vector ^56^ previously digested with HindIII and HpaI enzymes using an In-Fusion HD Cloning Kit (Takara-Clontech) following the manufacturer’s instructions. To express Myc-XPF fusion protein, the corresponding coding sequence was cloned at the NdeI and BamHI sites of pGRV23-myc. This plasmid is an expression vector which afford regulated expression of the corresponding gene from a procyclin promoter responsive to tetracycline and a puromycin N-acetyl- transferase resistance gene as selectable marker.

Ten micrograms of linear targeting DNA fragments obtained by NotI digestion were transfected into bloodstream parasites by electroporation using the Amaxa^TM^ Human T Cell Nucleofector Kit (Lonza) and the Amaxa Nucleofector device. Clones were selected with the appropriate selection drugs at the following concentrations: 0.1 µg/mL puromycin and 5 µg/mL hygromycin (Sigma). RNAi and *TbXPF* expression was induced using 1 µg/mL of doxycycline (Sigma). Selected clones of RNAi and TbXPF- expressing cells were analyzed by qPCR or Western blot analysis.

### Sensitivity assays

To determine the sensitivity of *T. brucei* cells lines to different genotoxic agents, a Resazurin Reduction Cell Viability Assay was performed in 96-well plates. Mid-log phase *T. brucei* parasites were diluted to a working cell density so that the final cell concentration was 75,000 cells/mL, and 90 µL/well was dispensed into 96-well flat-bottom transparent assay plates. Compounds at different concentrations were added to the cell plates (10 µL/well) which were incubated for 24 h at 37 °C and 5% CO_2_. Four hours prior to the end of the incubation, 20 µL of a Resazurin solution (440 µM, SIGMA) in pre-warmed HMI-9 was added to each well and incubated for another 4 h. Fluorescence was then measured in an Infinite F200 plate reader (TECAN infinite 200) at 550 nm (excitation filter) and 590 nm (emission filter). Assays were performed in duplicate at least twice to achieve a minimal *n*=2 per dose–response.

Camptothecin, phleomycin, mitomycin C, methyl methanesulfonate, hydrogen peroxide, cisplatin and doxorubicin were obtained from Merck Life Sciences. Methylnitronitrosoguanidine was purchased from MedchemExpress and fexinidazole from Selleck.

For UV irradiation, log-phase parasites at 5 x 10^3^ cells/mL in HMI-9 with 10% FBS were seeded in 60 mm Petri dishes prior to UVC irradiation (254 nm) at the indicated dose (UVP CL-1000 UV Crosslinker). After irradiation, plates were further incubated for 48 h at 37 °C before cells were counted using a Z1 Coulter counter.

### RNA extraction and real time quantitative PCR (RT-qPCR)

Total RNA was extracted from 5 x 10^7^ parasites using the NucleoSpin RNA kit (Macherey-Nagel) according to the manufacturer’s instructions. Reverse transcription was performed using iScript^™^ cDNA Synthesis kit (Bio-Rad) and quantitative PCR assays were carried out in a iCycler IQ real-time PCR detection system (Bio-Rad), with with 1X SYBR Green master mix (Thermo Scientific). Specific primers were designed to amplify *TbXPF* (Suppl. Table 1) and their efficiency was calculated from the equation E = 10^[-1/slope]^ obtained after plotting the cycle threshold (Ct) values *versus* the log of the cDNA amount ^57^. Relative expression of each gene was calculated with respect to the expression of a reference gene (*actin A*, Tb927.9.8850). Two independent experiments were performed and sample triplicates were used in all RT-qPCR assays.

### Subcellular localization studies

Intracellular localization of TbXPF was studied by immunofluorescence analysis. A total of 5 x 10^3^ cells were centrifuged and spread onto poly-L-lysine-coated slides. The cells were then fixed with 4% paraformaldehyde (diluted in 1x PBS) for 20 minutes at room temperature, washed twice, and permeabilized with 1% IGEPAL (Sigma-Aldrich) for 40 minutes. Next, the cells were blocked with 5% blocking reagent (Roche) for 30 minutes. The coverslips were first incubated with the primary antibody in blocking solution [(1X PBS, 0.5% blocking reagent (Roche)] for 1h and washed three times for 10 min. The primary antibodies used in this study were: mouse monoclonal anti-myc tag (clone 4A6) (1:200 dilution; Sigma-Aldrich); rabbit monoclonal anti-myc tag antibody (clone 71D10) (1:200 dilution; Cell Signalling Technology); mouse monoclonal anti-L1C6 (1:200 dilution; a gift from Dr. Bastin, Institut Pasteur, France).

Next, cells were incubated with Alexa Fluor 488 goat anti-mouse IgG or Alexa Fluor 594 goat anti-rabbit IgG secondary antibodies (Sigma, 1:1000 diluted in blocking solution) for 1 h. After washing, coverslips were dehydrated in methanol for 1 min and stained and mounted with Vectashield-DAPI (Vector Laboratories, Inc.). Vertical stacks of up to 40 slices (0.2 µm steps) were captured using an inverted Leica DMi8 microscope, 100x objective, and LASX software. Images were deconvolved and pseudo-colored with Huygens Essential software (version 3.3; Scientific Volume Imaging) and processed with ImageJ software (version 1.37; National Institutes of Health).

### Fluorescence-activated cell sorting (FACS) analysis

Parasites were harvested by centrifugation (1000 x *g*, 4 °C, 10 min) in mid log-phase (1 x 10 ^6^ cells/mL). Samples were washed twice in PBS, resuspended in 70% ice-cold ethanol/PBS solution and fixed overnight at 4 °C. Cells were stained and treated with 40 μg/mL of propidium iodide and 10 μg/mL of RNase A, respectively, in 500 μl of PBS during 30 min at room temperature in dark. FACS analysis of the cell cycle was carried out with a FACSCalibur flow cytometer (Becton Dickinson) and FlowJo software v10.0.

## ACKNOWLEDGMENTS

This work was funded by MCIN/AEI/ 10.13039/501100011033 and by “ERDF A way of making Europe”, by the “European Union” (Grants PID2021-124911OB-I00; PID2022-142971OB-I00).

## AUTHOR CONTRIBUTIONS

AEV designed the project and AEV and CGL, the experiments; CGL and MSM performed the experiments; CGL, MSM, LMRP, DGP and AEV analyzed the experiments; AEV wrote the manuscript; and CGL, MSM, LMRP, DGP and AEV edited and approved the manuscript.

## DATA AVAILABILITY

The authors declare that the data supporting the findings of this study are available within the article and its supplementary information files.

