## Supplemental Table 1 for "Role of the Nucleotide Excision Repair endonuclease XPF in the kinetoplastid parasite *Trypanosoma brucei*"

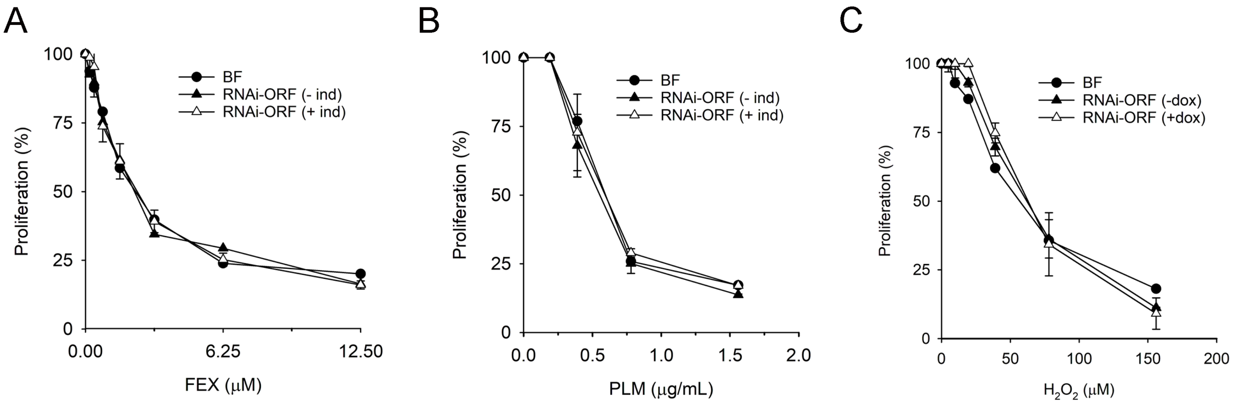


**Supplemental Figure 1. Proliferation assay of NER-defective and parental parasites exposed to genotoxic compounds.** Log-phase parasites at 5 x $10^3$ cells/mL were exposed to increasing concentrations of (A) fexinidazole (FEX); (B) phleomycin (PLM); and (C) hydrogen peroxide ($H_2O_2$) for 24 h at 37 ºC. Cell growth was measured with resazurin as described in Materials and Methods. For each cell line, proliferation was calculated relative to cell growth in the absence of compound. Experiments were performed at least three times, in duplicate. Values are the mean (±SD).

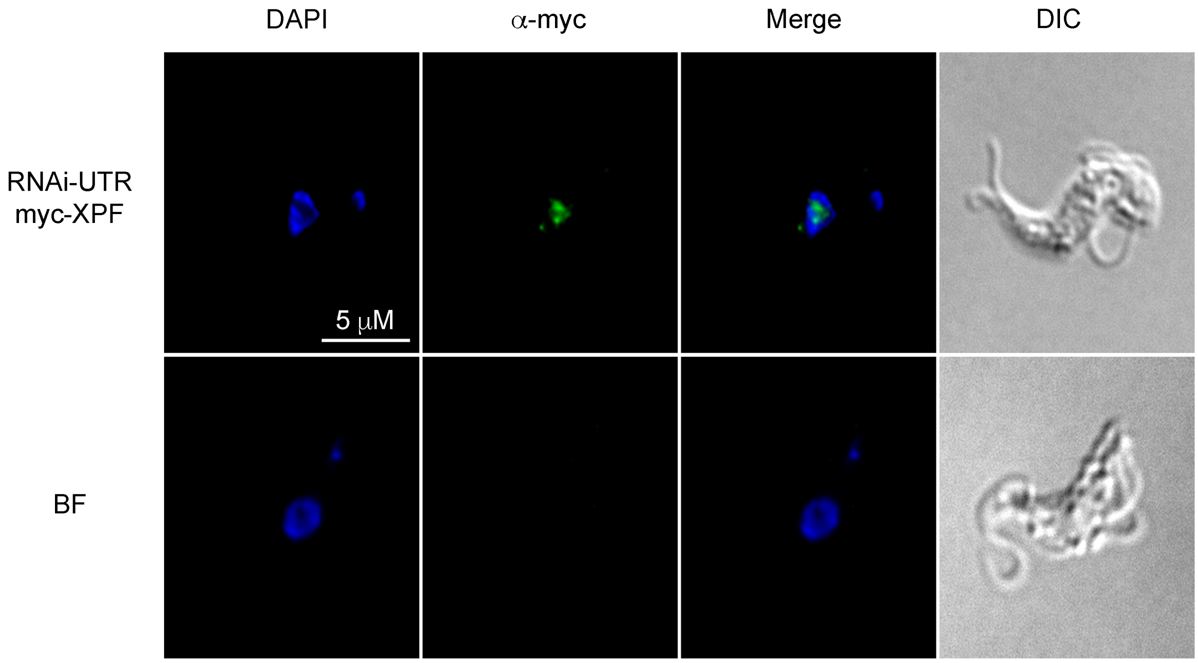


**Supplemental Figure 2. Detection of TbXPF in the nucleolar region using monoclonal anti-myc tag antibody (clone 4A6).** Immunofluorescence microscopy images from parental (BF) and myc-expressing parasites (RNAi-UTR myc-XPF) were obtained using an anti-myc tag mouse monoclonal antibody (clone 4A6) and Alexa Fluor 488 goat anti-mouse secondary antibody. Nuclear and kinetoplast DNA were stained with DAPI. Images were collected with an inverted Leica DMi8 microscope, 100x objective and LASX software.

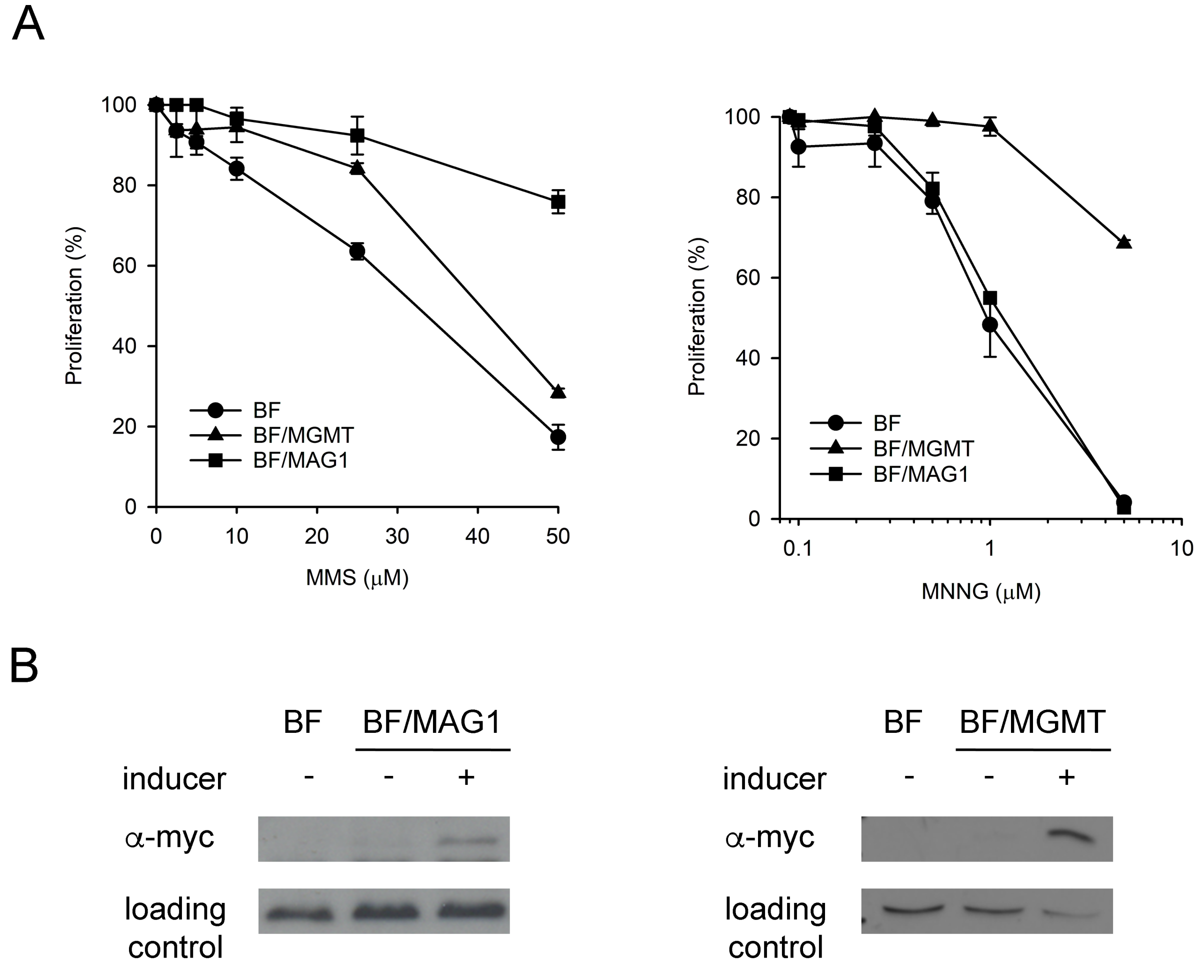


**Supplemental Figure 3. Assessing DNA damage induced by SN1 and SN2 methylating agents.** (A) Parental (BF) and transgenic bloodstream *T. brucei* cell lines expressing human MGMT protein (BF/MGMT) or the MAG1 protein from *Saccharomyces cerevisiae* (BF/MAG1) parasites (5 x $10^3$ cells/mL) were exposed to increasing concentrations of methyl methanesulfonate (MMS) and methyl-nitro-nitrosoguanidine (MNNG) for 48 h at 37 ºC. The curves show the mean of at least three experiments for each compound, values are the mean (±SD). (B) Western blot analysis showing the expression of human MGMT and yeast MAG1 in the bloodstream transgenic cell lines.

**Supplemental Table 1. List of oligonucleotides used in this study**.

| Sequence (5'>3') | Description | Use |
|---|---|---|
| AAA AAG TAA AAT TCA CAA GCT TGA TGG CAC AAG ATG TGT TAT C | Forward primer to amplify the ORF of XPF. Contains a complementary sequence to pGR19 and a HindIII site | Generation of an RNAi vector by In-fusion cloning |
| GAT GGC CGC TCT AGA ACT AGC TGC GGT CAA ACT CAA TAA G | Reverse primer to amplify the ORF of XPF. Contains a complementary sequence to the Stuffer fragment | Generation of an RNAi vector by In-fusion cloning |
| CTG GGG CGT GCA GGA CCA GCC GCT GCG GTC AAA CTC AAT AAG | Forward primer to amplify the ORF of XPF. Contains a complementary sequence to the Stuffer fragment. | Generation of an RNAi vector by In-fusion cloning |
| CAA CCC GGT GTT AGG ATC CGT TGA TGG CAC AAG ATG TGT TAT C | Reverse primer to amplify the ORF of XPF. Contains a complementary sequence to pGR19 | Generation of an RNAi vector by In-fusion cloning |
| CTA GTT CTA GAG CGG CCA TC | Forward primer to amplify the Stuffer fragment of pGR19 | Generation of an RNAi vector by In-fusion cloning |
| CGG CTG GTC CTG CAC GCC CCA G | Reverse primer to amplify the Stuffer fragment of pGR19 | Generation of an RNAi vector by In-fusion cloning |
| AAA AAG TAA AAT TCA CAA GCT TAC TTG TGT TTG GTT GTG TTA G | Forward primer to amplify the 3'UTR of XPF. Contains a complementary sequence to pGR19 and a HindIII site | Generation of an RNAi vector by In-fusion cloning |
| GAT GGC CGC TCT AGA ACT AGC ATA CCT CCA CTC AGA GAA G | Reverse primer to amplify the 3'UTR of XPF. Contains a complementary sequence to the Stuffer fragment | Generation of an RNAi vector by In-fusion cloning |
| CTG GGG CGT GCA GGA CCA GCC GCA TAC CTC CAC TCA GAG AAG | Forward primer to amplify the 3'UTR of XPF. Contains a complementary sequence to the Stuffer fragment. | Generation of an RNAi vector by In-fusion cloning |
| CAA CCC GGT GTT AGG ATC CGT TAC TTG TGT TTG GTT GTG TTA G | Reverse primer to amplify the 3'-UTR of XPF. Contains a complementary sequence to pGR19. | Generation of an RNAi vector by In-fusion cloning |
| GAT GGT TGT TGT GTT TGG TC | Forward primer (qPCR) | Quantification of TbXPF mRNA in RNAi-ORF cells |
| CTG CTT CTC GGT ATC GTT GTC | Reverse primer (qPCR) | Quantification of TbXPF mRNA in RNAi-ORF cells |
| CAG CGT GTC ATG GCA CTT TG | Forward primer (qPCR) | Quantification of TbXPF mRNA in RNAi-UTR cells |
| CTACATATTGGGTCTAACAC | Reverse primer (qPCR) | Quantification of TbXPF mRNA in RNAi-UTR cells |
| CAT ATG CCA CAG ACT GTT AGC | Forward primer to amplify TbXPF. Contains an NdeI site for cloning into pGRV23 | Expression of TbXPF in *T. brucei* cells |
| AGA TCT TTA TGT CTG AGT AGG TGG C | Reverse primer to amplify TbXPF. Contains a BglII site for cloning into pGRV23-myc | Expression of Myc-TbXPF in *T. brucei* cells |
| AGA TCT TTA TTT GTC GTC ATC GTC TTT GTA GTC TGT CTG AGT AGG TGG CAC | Reverse primer to amplify TbXPF. Includes a C-terminal FLAG tag. Contains a BglII site for cloning into pGRV23 | Expression of TbXPF-Flag in *T. brucei* cells |
| GAC CAT ATG GAC AAG GAT TGT GAA ATG | Forward oligo to amplify MGMT. Contains an NdeI site for cloning into pGRV23-myc | Expression of myc-MGMT in *T. brucei* cells |
| AGT GGA TCC TCA GTT TCG GCC AGC AGG C | Reverse oligo to amplify MGMT. Contains a BamHI site for cloning into pGRV23-myc | Expression of myc-MGMT in *T. brucei* cells |
| AGG GCA ATT AAT ATG GAG GAG CAG AAG CTG ATC | Forward oligo to amplify MAG1. Contains an AseI site for cloning into pGRV23-myc | Expression of myc-MAG1 in *T. brucei* cells |
| AAG CTT GGA TCC TTA GGA TTT CAC GAA ATT TTC | Reverse oligo to amplify MAG1. Contains a BamHI site for cloning into pGRV23-myc | Expression of myc-MAG1 in *T. brucei* cells |
